# Investigating the use of novel blood processing methods to boost the identification of biomarkers of non-small cell lung cancer

**DOI:** 10.1101/2024.09.18.613811

**Authors:** Rosalee McMahon, Natasha Lucas, Cameron Hill, Dana Pascovici, Ben Herbert, Elisabeth Karsten

## Abstract

1.

**Background and objectives:** Diagnosis of non-small cell lung cancer (NSCLC) currently relies on imaging and in-clinic visits, however these methods are not effective at detecting early-stage disease. The investigation into blood-based biomarkers aims to simplify the diagnostic process and has the potential for identifying disease-associated changes before they can be seen using imaging techniques.

**Methods and design:** In this study, plasma and frozen whole blood cell pellets from patients with NSCLC and healthy controls were processed using both classical as well as novel techniques to produce a unique set of 4 sample types from a single blood draw. Samples were analysed using 12 commercially available immunoassay kits in addition to liquid chromatography mass spectrometry using a Q Exactive HF-X Orbitrap to collectively screen 3974 proteins as potential biomarkers.

**Results and conclusions:** Analysis of all sample types produced a set of 522 differentially expressed proteins, with conventional blood analysis (proteomic analysis of plasma) accounting for only 7 of that total. Boosted regression tree analysis of the differentially expressed proteins produced a panel of 13 proteins that were able to discriminate between controls and NSCLC patients with an area under the ROC curve (AUC) of 0.864 for the set. Our rapid and reproducible blood preparation and analysis methods enable the production of high-quality data from small aliquots of complex samples that are typically seen as requiring significant fractionation prior to proteomic analysis.

## 2. Introduction

Biomarkers of disease, particularly those detectable in easily accessed biological fluids, are the hallmark of modern diagnostics. For example, prostate-specific antigen (PSA) is a highly effective biomarker for prostate cancer and is used as a first pass diagnostic in addition to monitoring disease progression.^1^ Despite a lack of specificity, analysis of this marker is used broadly and frequently, due to the benefits that come with a marker that is easily identifiable in a blood sample and not reliant on imaging or more onerous protocols.

No equivalent biomarker exists for lung cancer, let alone for non-small cell lung carcinoma (NSCLC).^2^ Diagnostics and disease monitoring are entirely reliant on imaging, typically X-rays followed by a CT-scan. Although specific, they lack sensitivity and are not used for broad, early-stage population screening. As such, NSCLC is typically left undiagnosed until Stage III or IV resulting in a 5-year survival rate of less than 15 %.^3^ Thus, the search is on for blood-based markers that may be diagnostic or even prognostic for NSCLC. The most promising biomarkers currently known (CEA and CYFRA21-1) lack the sensitivity and specificity required for clinic adoption.^4^ Investigations have since deduced that a single biomarker is unlikely to be sufficient for NSCLC diagnosis, instead combinations of multiple markers may be more promising.^5,6^

Although there is a lot known about the role of leukocytes in cancer progression, it is now understood that red blood cells (RBCs) may too play a role in cell-cell communication.^7^ Our laboratory found that a number of key inflammatory markers are stored at high levels in these cells and that not all of the markers are released and available for analysis in plasma or serum.^8,9^ *In vitro* analysis revealed that by co-culturing RBCs with a NSCLC cell line, the signalling molecules and protein markers within RBCs can change substantially.^10^ In a model of prostate cancer, a chemokine receptor on RBCs was found to be critical in regulating tumour growth and in reducing intra-tumoural chemokine levels.^11^ These studies indicate that, *in vivo*, a tumour burden is likely to influence the protein profile of RBCs, making them a potentially untapped source of unique disease biomarkers.

One sampling method that, by design, produces a sample that includes RBCs are dried blood spots and volumetric absorptive microsampling (VAMS) devices. These are typically marketed as at-home whole blood (WB) sampling devices. This is due to the ease of sample collection wherein only small volumes are required and can be collected from a finger prick and the samples are not classified as a dangerous good so can be shipped in the regular mail once dried.^12^ We have discovered an additional utility of these samples as sample preparation devices.^13^ Discovery proteomics of blood samples using mass spectrometry is limited due to the significant dynamic range of proteins in blood. By using dried blood spots or VAMS as a sample preparation device, a subset of proteins from blood are immobilised and with specific wash buffers, high abundance proteins can be removed, leaving behind a set of potentially interesting proteins. These methods are highly reproducible and can be modified to select for particular protein subsets of interest.^13^ This enables more protein identifications from a whole blood sample than could be achieved with corresponding liquid blood.

The aim of this study was, using multi-fractionated blood analysis, to identify a unique set of differentially expressed proteins for NSCLC that have not been previously identified with conventional plasma or serum proteomics.

## 3. Materials and Methods

### 3.1. Materials

Na2HPO4, NaH2PO4, LiCl, Tris, TCEP, chloroacetamide, triethylammonium bicarbonate buffer, sodium deoxycholate, formic acid (FA) and acetonitrile were purchased from Sigma Aldrich (Castle Hill, Australia) and were analytical grade. Phosphate buffered saline (PBS) and Pierce BCA (bicinchoninic acid) protein assay were purchased from Thermo Fisher (Scoresby, Australia). Roche cOmplete protease inhibitor cocktail was purchased from Sigma Aldrich (Castle Hill, Australia). Trypsin Gold was of mass spectrometry grade and was purchased from Promega (Alexandria, Australia). Oasis HLB 1cc cartridges (10mg sorbent, 30 μm) were purchased from Waters (Rydalmere, Australia). 30 μL Mitra volumetric absorptive microsampling devices were purchased from Trajan (Neoteryx, Ringwood, Australia). All blood samples were purchased from Precision Med, Inc (Solana Beach, CA, USA). Blood samples consisted of EDTA plasma and corresponding cell pellets from 16 NSCLC patients and 18 age and sex matched controls. Cell pellets were stored frozen at -20 °C with the supplier and were transferred to -80 °C upon delivery. Plasma was stored frozen at -80 °C with the supplier and upon receipt in the laboratory. Immunoassay were purchased from various manufacturers outlined in **Table 1**.

**Table 1.** Summary of immunoassays used to quantify proteins in blood samples.

| Kits | Manufacturer | Analytes assayed |
| --- | --- | --- |
| **Luminex®** |  |  |
| Human high sensitivity T-cell panel, 21-plex, Milliplex | Millipore | Fractalkine, GM-CSF, IFNγ, IL-1β, IL-2, IL-4, IL-5, IL-6, IL-7, IL-8, IL-10, IL-12, IL-13, IL-17A, IL-21, IL-23, ITAC, MIP-1α, MIP-1β, MIP-3α, TNFα |
| Custom chemokine panel, 12-plex, ProcartaPlex (Invitrogen) | Invitrogen | ENA-78, Eotaxin, GRO-α, IL-8, IP-10, ITAC, MCP-1, MIG, MIP-1α, MIP-1β, MIP-3α, RANTES |
| Human angiogenesis panel 2, 20-plex, MilliPlex (Millipore) | Millipore | Angiostatin, Osteopontin, PDGF-AA/BB, sAXL, sc-Kit/SCFR, sEGFR, sE-selectin, sHer2, sHer3, sHGFR/c-Met, sIL-6Ra, sNeuropilin-1, sPECAM-1, sTie-2, sVEGFR1, sVEGFR2, sVEGFR3, suPAR, Tenascin C, Thrombospondin-2 |
| MIF single-plex (R&D Systems) | R&D Systems | MIF |
| **ELISAs** |  |  |
| IL-8 | Elisakit | IL-8 |
| ITAC | Abcam | ITAC |
| MCP-1 | Abcam | MCP-1 |
| Midkine | Abcam | Midkine |
| MIF | Abcam | MIF |
| MIP-1α | Abcam | MIP-1α |
| TNF-α | Thermo Fisher | TNF-α |
| **Flow cytometry bead-based array** |  |  |
| Growth factor, 13-plex panel, LegendPlex | BioLegend | Angiopoietin-2, EGF, EPO, FGFb, G-CSF, GM-CSF, HGF, M-CSF, PDGF-AA, PDGF-BB, SCF, TGF-α, VEGF |

Sodium phosphate buffer (5mM, pH 7.4) was prepared by reconstituting Na2HPO4 (66% w/v) and NaH2PO4 (8% w/v) in Milli-Q water. Extraction buffer was comprised of 500mM LiCl + 100 mM Tris. Lysis buffer (pH 8) was comprised of 1 % sodium-deoxycholate (w/v), 10 mM TCEP, 40 mM chloroacetamide, and 100 mM triethylammonium bicarbonate buffer.

### 3.2. Immunoassays

#### 3.2.1. Blood sample preparation

The whole blood cell pellet samples were processed to produce two fractions, (1) a membrane-free supernatant fraction (soluble proteins released after erythrocyte lysis) and (2) an erythrocyte membrane release fraction (proteins released during overnight incubation form washed erythrocyte membranes). These fractions were prepared as follows.

The cell pellets were homogenised and diluted 12.5x to 1500 μL with sodium phosphate buffer and left to incubate (5 mins, room temperature). The sample were vortexed to encourage cell lysis and then centrifuged (2,000 *g*, 5 mins, room temperature). The supernatant was collected and was further centrifuged to isolate the erythrocyte membranes (20,000 *g*, 20 mins, 4 °C). The resulting membrane-free supernatant was collected and frozen at -80 °C in aliquots until analysis.

The remaining membrane pellet was washed three times in sodium phosphate buffer followed by a single wash in PBS (20,000 *g*, 20 mins, 4 °C). The membranes were then resuspended in 1000 μL PBS with protease inhibitors and incubated (24 hrs, 37 °C). Following incubation, the membrane suspensions were centrifuged (20,000 *g*, 20 mins, 4 °C) and the membrane releasates were collected at -80 °C in aliquots until analysis.

#### 3.2.2. Protein quantification

All blood fractions were analysed using Luminex® protein detection panels, ELISA kits, and a flow cytometry bead array (LegendPlex™). Details of the specific assays are summarised in **Supplementary Table 1**. Kits were run according to manufacturers instructions with samples dilutions optimised for each assay. Luminex® assays were run on the Luminex 200 system (Bio-Rad). The calibration curve for each cytokine was analysis with 5-parametric logistic curve regression using Bio-Plex manager software (ver. 5.0, Bio-Rad). ELISAs were performed with quantification of absorbance using a spectrophotometer (BioTek). Calibration curves were prepared as recommended by the manufacturer. The flow cytometry bead array was performed with quantification of proteins using LSRFortessa flow cytometer (BD) to collect 5000 events per sample. Calibration curve and data analysis was performed using LegendPlex analysis software (ver. 8.0, BioLegend).

#### 3.2.3. Sample normalisation

Data was normalised across participants according to haemoglobin concentration for the membrane-free fraction and total protein levels for membrane releasates. Levels of haemoglobin were quantified by assessing absorbance at 414 nm (Synergy 2 Multi-Mode plate reader, BioTek). Membrane-free samples were normalised using the results of a single participant as the control.

Total protein levels of the membrane releasates were determined by BCA assay according to manufacturers instructions. The immunoassay results for the membrane releasate samples were normalised to 1 mg/mL of total protein.

### 3.3. Mass spectrometry

#### 3.3.1. Sample preparation

Samples were randomised and processed in 2 batches. The WB cell pellets were thawed and diluted 1:1 in PBS. VAMS were prepared by dipping Mitra® microsampling tips (30 μL) into the diluted WB cell pellets until the tip had wicked up the blood and was full. A single 30 μL tip was prepared for each sample. All samples were then air-dried for 5 minutes at room temperature before being transferred to a foil ziplock bag with desiccant and were left to completely dry for 24 hours at room temperature.

The samples were then extracted immediately. The tips were removed from the plastic spindles and were suspended in 1 mL extraction solution and incubated (24 hrs, RT) whilst gently shaking. The tips were then washed twice in extraction solution. Proteins were digested by suspending the washed tips in 100 μL lysis buffer and were heated at 95 °C for 10 mins with agitation (Thermomixer, Eppendorf). After heating, 1 μg trypsin was added (1 μg/μL in lysis buffer) and the sample was overnight at (19 hours, 37 °C). Samples were then diluted with 800 μL of 0.5 % formic acid and the tip was removed from the liquid sample and discarded. The remaining sample was centrifuged (16,000 *g*, 10 mins) and the supernatant was transferred to a conditioned solid phase extraction cartridge (Oasis HLB cartridge). Sample was desalted using solid phase extraction. Peptide concentration was determined using Nanodrop spectrophotometer (Thermo Fisher).

#### 3.3.2. Mass spectrometry acquisition

All samples were acquired in duplicate. Peptides (0.5 μg) were loaded directly onto a nano liquid chromatography column (75 μm x 50 cm 1.9 μm C18 ReproSil-Pur 120 C18-AQ) in 97 % solvent A (0.1 % FA) and 3 % solvent B (80 % acetonitrile, 0.1 % FA) at 500 μL/min for 12 mins. The flow rate was decreased to 300 μL/min and peptides were eluted over 62.5 min gradient to a maximum of 30 % solvent B. The gradient was then stepped up to 60 % solvent B over 2.5 min, before washing in 98 % solvent B for 5 mins.

Peptides were detected using a HF-X Orbitrap mass spectrometer (Thermo, CA, USA). Data was acquired in data-independent acquisition (DIA) mode using the following instrument parameters: Full MS scan resolution 120 K, AGC 3 x 10^6^, maximum injection time 50 ms, scan range 350-1650 m/z, followed by 20 DIA variable windows to scan 350-1650 m/z as per **Supplementary Table S1** with settings of Resolution 30 K, AGC 3 x 10^6^, maximum injection time set to auto. The collision energy was stepped at 22.5, 25, 27.5, default state was 3.

#### 3.3.3. Mass spectrometry controls

A HeLa digest was run before each batch of samples were acquired on the mass spectrometer. A total of 5 runs were acquired.

#### 3.3.4. DIA-NN analysis

All 64 files were processed through DIA-NN software using the following settings: Output filtered at 0.01 FDR; Precursor/protein x samples expression level matrices saved along with the main report; Deep learning used to generate a new in silico spectral library from peptides provided; Library-free search enabled; Min fragment m/z set to 200; Max fragment m/z set to 1800; N-terminal methionine excision enabled; In silico digest will involve cuts at K*,R*; Maximum number of missed cleavages set to 2; Min peptide length set to 7; Max peptide length set to 30; Min precursor m/z to 300; Max precursor m/z set to 1800; Min precursor charge set to 2; Max precursor charge set to 5; Cysteine carbamidomethylation enabled as a fixed modification; A spectral library will be created from the DIA runs and used to re-analyse them. DIA-NN will optimise the mass accuracy automatically using the first run in the experiment.

### 3.4. Data analysis

Data analysis and graphing of data was performed using GraphPad Prism (ver. 8.0) and SPSS (ver. 26). Comparison between healthy and NSCLC patients was performed using 2-tailed, unpaired, Student t-tests. ANCOVA analysis was also performed to correct for covariates (sample age (time in storage) and sex), cytokine concentration as the dependent variable, and cancer status (healthy vs NSCLC) as the independent variable. Analysis of cytokine concentration per cancer stage (healthy, Stage II, Stage III, or Stage IV) was determined using 1-way ANOVA. Spearman correlations between cytokine concentration and sample age, sex, or clinical stage were performed and graphed using GraphPad Prism (ver. 8.0). Data was deemed statistically significant if p < 0.05.

Mass spectrometry data was processed and analysed using R Studio (ver. 2022.02.1). Data was filtered to remove sparse proteins with fewer than 3 quantitated values in both groups. Remaining protein quantitation with zero or missing values were imputed using the default Perseus-style imputation, wherein random numbers were drawn from a normal distribution of 1.8 standard deviation down shift and with a width of 0.3 of each sample. Median normalisation was carried out before data imputation and no batch normalisation was undertaken. After data imputation, differential expression analysis was carried out. Initially the data was compared between groups with a two-sample t-test and proteins were regarded as differentially expressed (DE) if the false discovery rate corrected p-vale was less than 0.5. Ingenuity Pathway Analysis (Ingenuity Systems Inc) was applied to describe protein-protein interaction networks and to consider the biological implications of differential expression of proteins.

A separate filter was undertaken to identify a smaller subset of proteins of interest. The filter calculated the importance rank of each protein using boosted regression methods as implemented in the R package GBM.^14,15^ The following key parameters were used for this analysis: the number of regression trees (n = 100), a Bernoulli distribution, a shrinkage parameter of 0.01, five-fold crossvalidation and a fraction of training data of 75 %. In each case, the top 10 ranked markers in terms of importance were retained. The procedure was repeated 100 times, and the markers were ranked in terms of the number of times they were selected in the top 10 importance rank for each protein. This ranking was combined with selected DE proteins based on a linear model including both cancer status and sex effect, where however there is no significant effect of sex, and no significant interaction between cancer status and sex.

## 4. Results

Blood samples were fractionated into 4 unique and complimentary samples (plasma, erythrocyte membrane-free WB cell lysate, erythrocyte membrane releasate, tip digest) with the aim to produce a diverse dataset for biomarker discovery. These samples were analysed using a variety of methods including Luminex, ELISA, flow cytometry, and mass spectrometry.

### 4.1. Demographics

Healthy controls (n = 18) were age and sex matched to the NSCLC participants (n = 16). A summary of participant demographic information and disease status is outline in **Table 2**. Clinical stage of NSCLC patients varied between II – IV. The average age of all participants was 65.3 ± 9.0 years and 81.3 % were of Caucasian ethnicity.

**Table 2.**
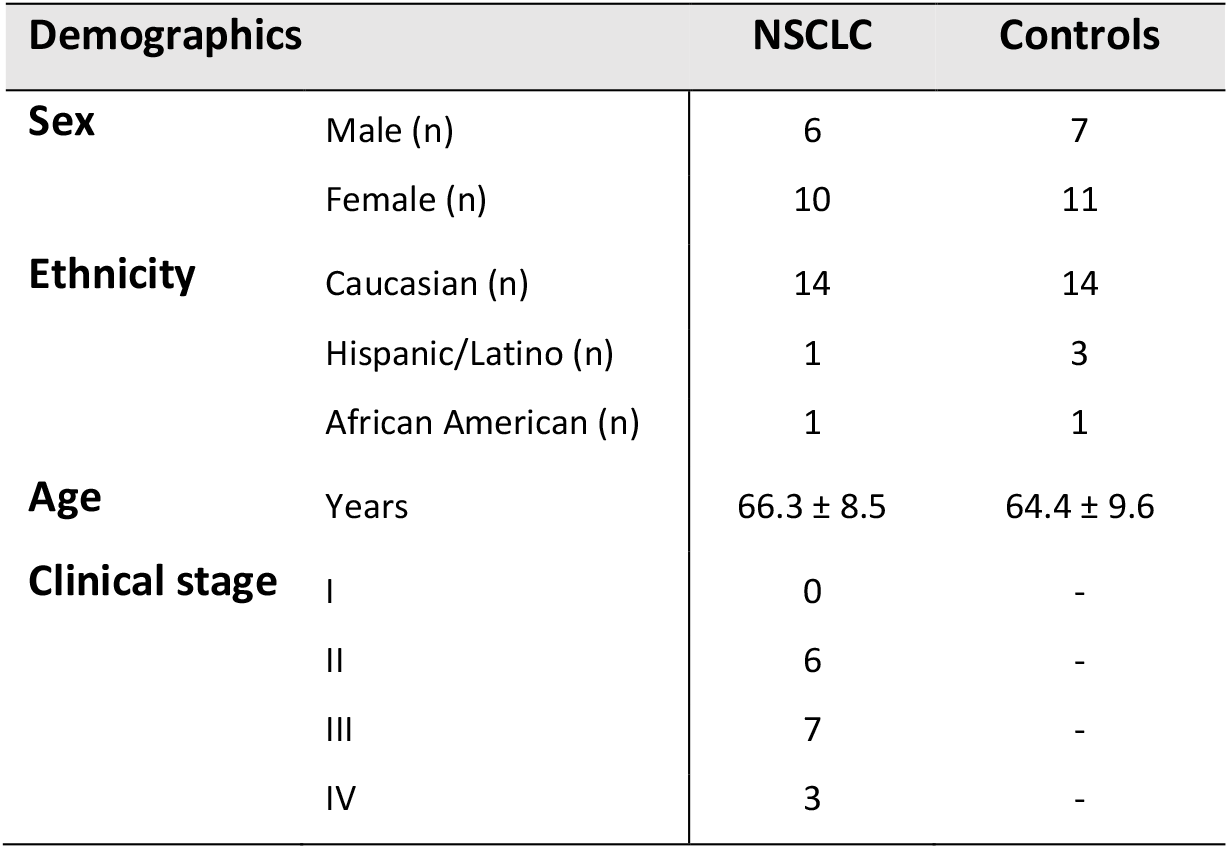
Summary of participant demographics and disease status.

| Demographics |  | NSCLC | Controls |
| --- | --- | --- | --- |
| Sex | Male (n) | 6 | 7 |
|  | Female (n) | 10 | 11 |
| Ethnicity | Caucasian (n) | 14 | 14 |
|  | Hispanic/Latino (n) | 1 | 3 |
|  | African American (n) | 1 | 1 |
| Age | Years | 66.3 ± 8.5 | 64.4 ± 9.6 |
| Clinical stage | I | 0 | - |
|  | II | 6 | - |
|  | III | 7 | - |
|  | IV | 3 | - |

### 4.2. Cytokine analysis

#### 4.2.1. Sample age

The biospecimen were collected over a period of 5 years prior to purchase and there was a disparity in the age of the samples between healthy participants and NSCLC participants. The age of samples from healthy participants ranged from 35.6 - 263.0 weeks and 43.0 - 165.4 weeks for NSCLC participants. Thus, it was of interest to determine if sample storage time had any effect on detected analyte concentration as this would create bias. Spearman correlation analysis revealed that 6 analytes had a significant correlation of r > 0.5 with sample age (**Supplementary Figure S1**). All were positive correlations and 5 out 6 were detected in the membrane-free fraction samples.

Prior to purchase the cell pellet from the blood tubes had been stored at -20 °C whilst the plasma fraction was stored at -80 °C. After the samples were delivered they were processed and stored at - 80 °C until analysis, however the initial difference in storage is likely to have contributed to these results. To account for these differences, age of sample was included as a covariate in the ANCOVA analysis of healthy vs NSCLC participants (**Figure 1**).

**Figure 1.**
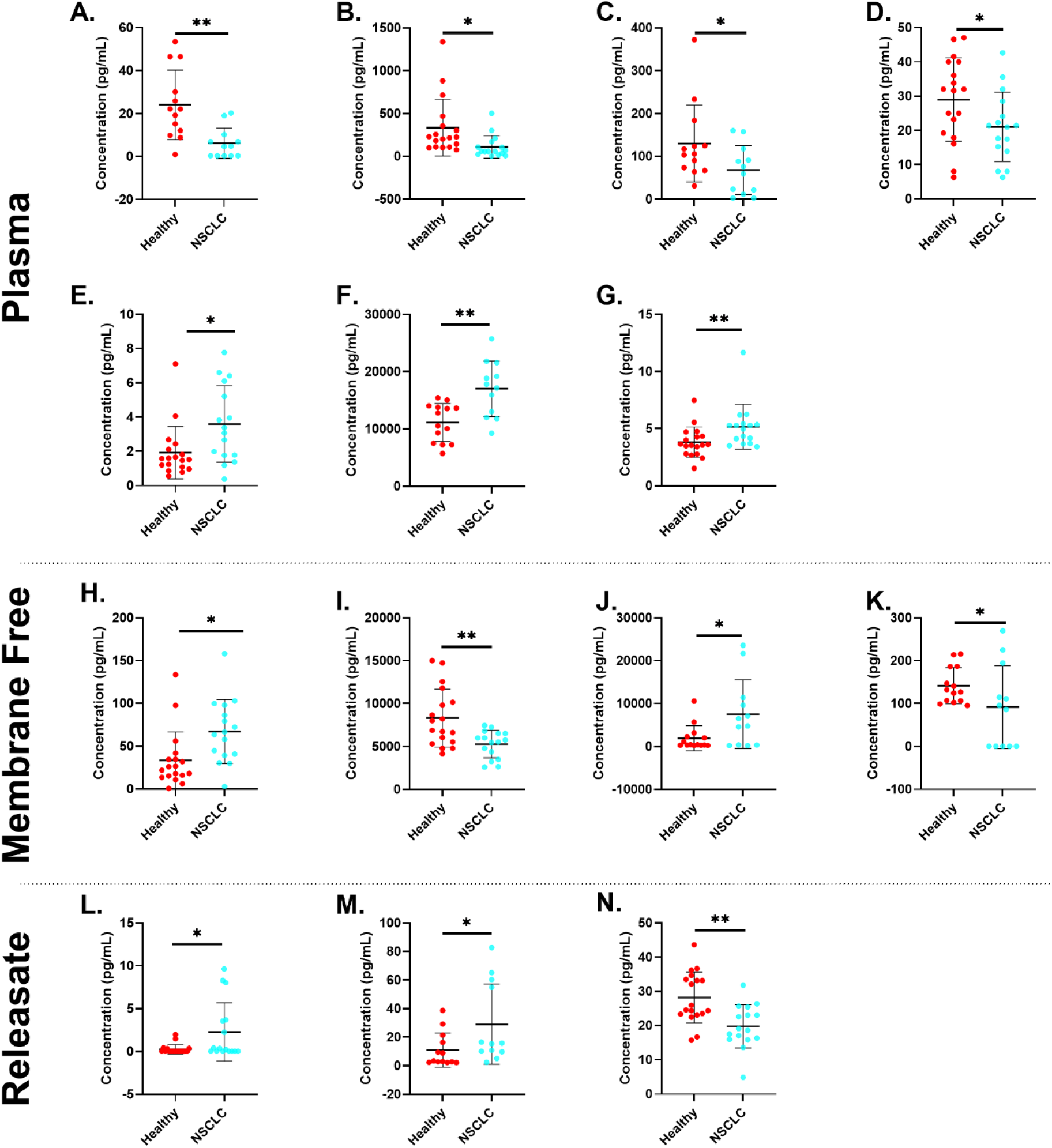
Concentration of (A) EGF, (B) ENA-78, (C) GCS-F, (D) IL-4, (E) IL-8, (F) Tenascin C and (G) TNF-a in plasma; (H) IL-8, (I) MIF, (J) suPAR, (K) sVEGFR2 detected in membrane-free fraction of whole blood cell pellet; and (L) GM-CSF, (M) HGF, MIP-1α in membrane releasate samples. Blood fractions were collected from healthy controls (red) and participants with NSCLC (blue) and measured by Luminex, ELISAs, or flow cytometry bead arrays. Data identified as statistically significant following ANCOVA analysis with correction for sample age and sex. Data presented as mean ± SD and deemed significant if p<0.05.

#### 4.2.2. Differential expression

Blood collected from healthy controls and NSCLC participants was fractionated and analysed using a variety of Luminex assays, ELISAs, and a flow cytometry bead array resulting in a total of 60 individually analysed proteins. ANCOVA analysis was performed to correct for sample age and sex differences, revealing 13 statistically significant proteins. In total, 7 were identified in the plasma fraction (EGF, ENA-78, G-CSF, IL-4, IL-8, Tenascin C, and TNFα), 3 in the membrane-free fraction (GM-CSF, HGF and MIP-1α), and 4 in the membrane releasate fraction (IL-8, MIF, sVEGFR2, and suPAR). Graphical summary of statistically significant results can be found in **Figure 1**.

The NSCLC participants in this study had varying stages of the disease (stage II – IV) when the biospecimen were collected. Any stage specific cytokine changes or overall trend in cytokine concentration with worsening disease was investigated with Spearman correlations. This analysis revealed 4 analytes with a significant correlation of > 0.7 or < -0.7. Two of these were detected in the plasma fraction (sVEGFR1 & sVEGFR2) and two in the membrane-free fraction (sc-Kit/SCFR & sNeuropillin-1). The strongest correlation identified was for sc-Kit/SCFR in the membrane-free fraction. There was an increase in levels of sc-Kit/SCFR with increasing clinical grade with a correlation of r = 0.825.

Although the sample size of each clinical stage groups is small (Stage II: n = 6; Stage III: n = 7; Stage IV: n = 3), there were some notable differences identified. For example, in the comparison between healthy and NSCLC participants, MIP-1α in the membrane-free fraction was identified as significantly different (**Figure 1**), however from the analysis of the individual cohorts, this difference appears to be driven by the participants with clinical stage III NSCLC (**Figure 2**). There is no significant difference between the healthy cohort and stage II or stage IV, however in stage III there is a significant decrease in the protein concentration. Likewise, there is a significant increase in sc-Kit/SCFR in the plasma in stage III participants compared to stage II. EGF levels in plasma are at their highest in healthy participants and are reduced in every stage of the disease that was analysed.

**Figure 2.**
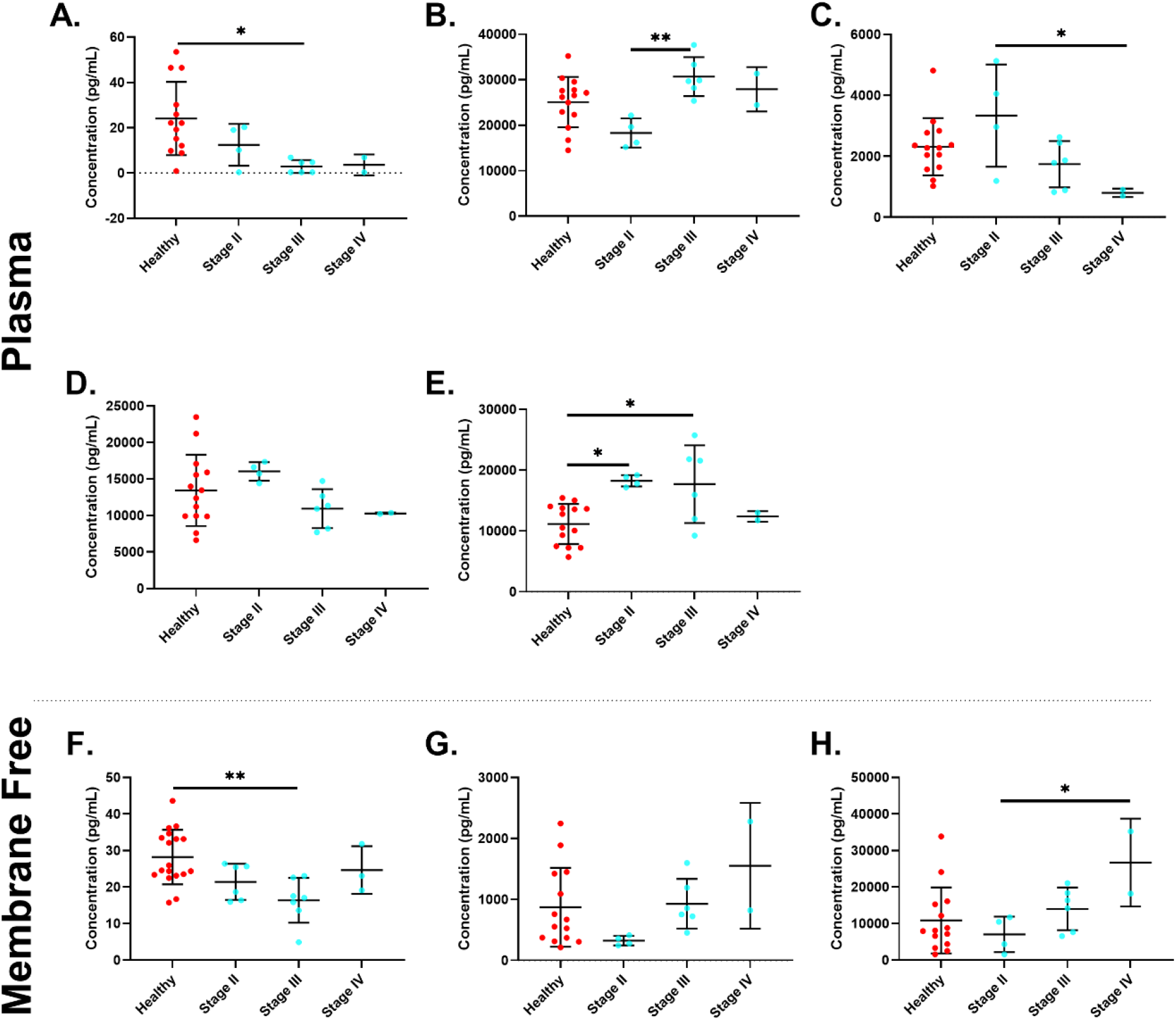
Concentration of (A) EGF, (B) sc-Kit/SCFR, (C) sVEGFR1, (D) sVEGFR2, (E) Tenascin C in plasma; and (F) MIP-1α, (G) sc-Kit/SCFR, (H) sNeuropilin-1 in membrane-free fraction of whole blood cell pellet collected from healthy participants and participants with NSCLC stratified by clinical stage and as measured by Luminex, ELISAs, or flow cytometry bead arrays. Data identified as statistically significant following ANOVA analysis. Data presented as mean ± SD and deemed significant if p < 0.05.

Conversely, sVEGFR2 (plasma) and sc-Kit/SCRF (membrane-free) were both found to have a significant correlation across clinical stage (r = -0.75 and r = 0.83 respectively) but no statistically significant differences were found between the individual groups (**Figure 2**).

### 4.3. Mass spectrometry analysis

#### 4.3.1. Sample validity

Total protein yield from samples was comparable across both processing batches with a mean of 17.2 ± 5.9 μg and 19.0 ± 5.4 μg for each batch respectively. There was also good reproducibility between technical replicates, as illustrated by PCA (**Figure 3**). The mass spectrometry results were particularly unaffected by sample age, possibly due to the fact that samples are digested prior to analysis, so fragments may still be identifiable. This was contrary to the immunoassay results, likely since those assays require intact antibody binding domains, making them more affected by sample degradation (**Figure 3**).

**Figure 3.**
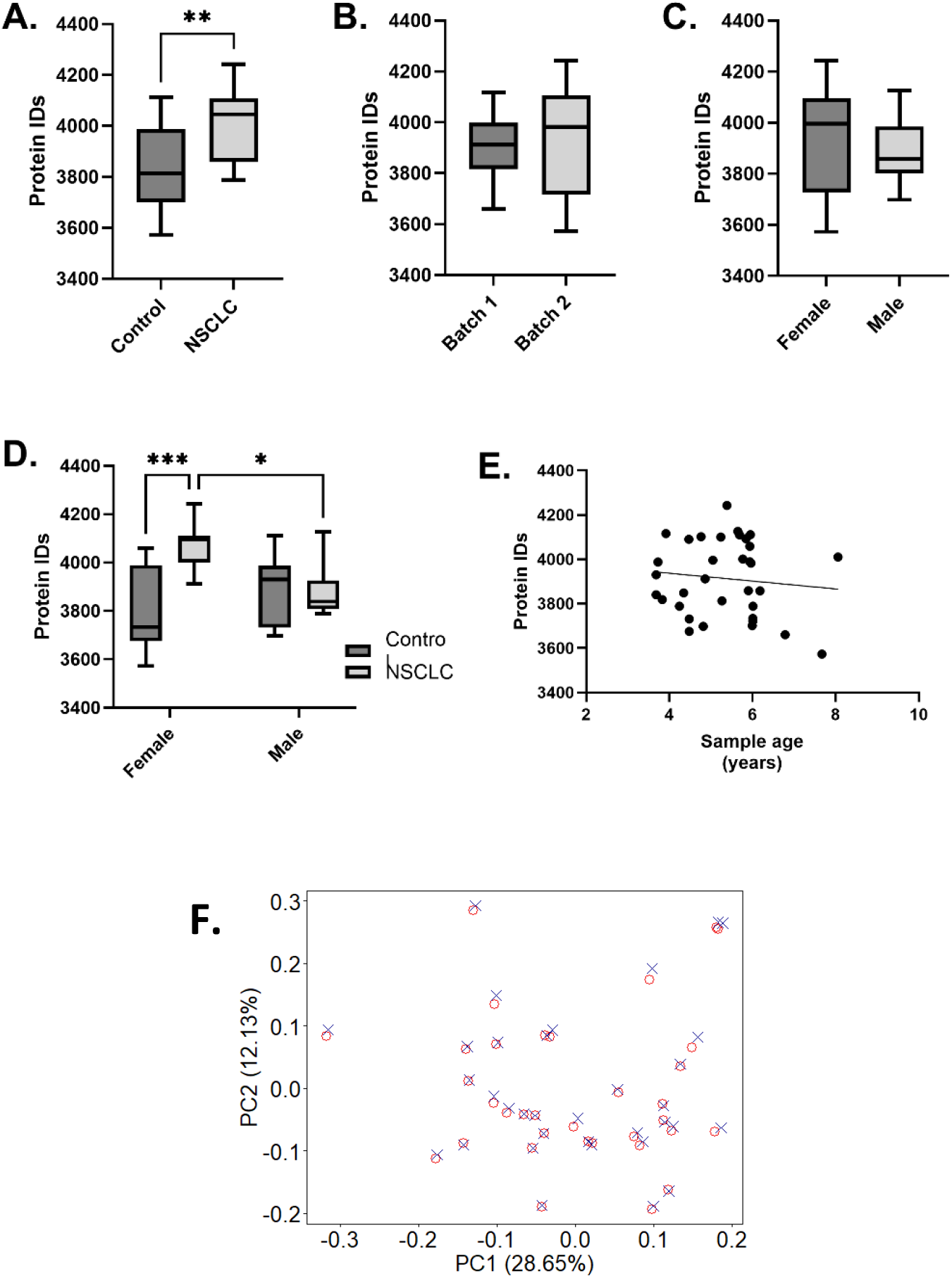
Number of protein IDs from blood samples stratified according to A) disease status, B) processing batch, C) sex, and D) sex and disease status combined, E) correlation between sample age (years) and protein IDs, and F) variation between technical replicates as assessed by PCA. Replicates are illustrated by × and ◯ respectively. Data is significant (*) if p < 0.05.

A mean of 3913.8 ± 170.9 proteins were identified across all samples with significantly more proteins identified in the NSCLC samples compared to the controls (4002 ± 141 and 3836 ± 160 protein IDs for NSCLC and controls respectively) (**Figure 3**). There was also a significant interaction observed for sex, wherein the NSCLC female cohort has significantly higher protein IDs than the male and control counterparts (**Figure 3**). This is likely driving the overall increase in IDs in the whole NSCLC cohort. No interaction with processing batches or correlation with age of sample was observed.

#### 4.3.2. Sex effect

There was a subset of female cancer samples that were well distinguished from the rest (**Figure 4**). When stratified for sex, 1034 differentially expressed (DE) proteins were identified for the female cohort and 0 DE proteins identified for the male cohort. In the DE proteins for the female cohort, 83.0 % were increased in NSCLC with an average log fold change of 1.3 ± 0.6. Restriction of DE proteins for the female cohort to those with p <0.05 and a fold change of > 1 or < -1 produced a list of 473 DE proteins. Ingenuity pathway analysis (IPA) of this subset revealed a canonical sirtuin signalling pathway that is decreased in the female NSCLC cohort, notably, this pathway is absent in IPA when not stratified by sex.

**Figure 4.**
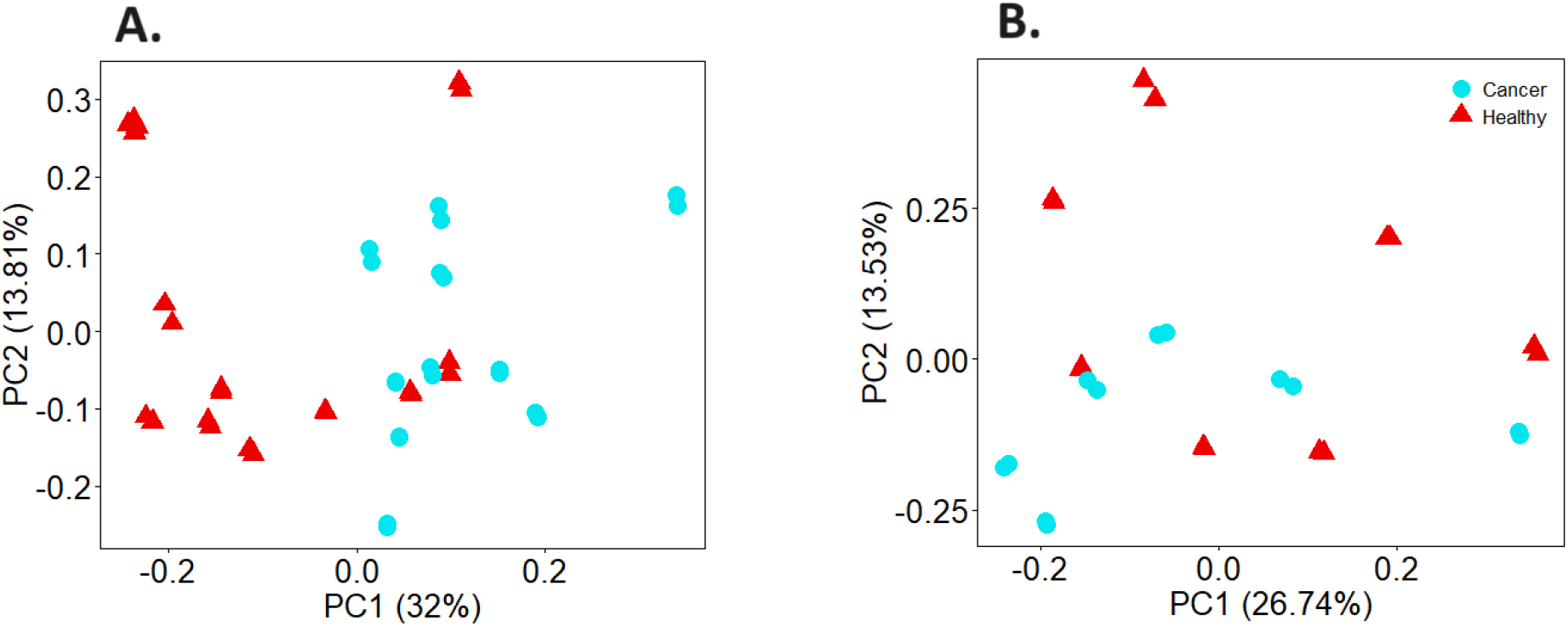
PCA plots of NSCLC patients (•) compared to controls (▴) separated according to sex wherein (A) are females (n = 10 and 11 for NSCLC and controls respectively) and (B) are males (n = 7 and n = 6 for NSCLC and controls respectively). All technical replicates included in each figure.

#### 4.3.3. Differential expression

Using standard criteria (false discovery rate adjusted p-values < 0.05), 508 DE proteins were identified. Ingenuity pathway analysis (IPA) revealed these identified proteins are involved with a variety of functional pathways covering adhesion and migration, as well as a strong network of known cancer-associated cytokines and enzymes including TGFB1 and IL1B (**Figure 5**). Further pathway analysis revealed that 39 were known biomarkers of cancer including CEA cell adhesion proteins 1 and 6, from the CEA family.

**Figure 5.**
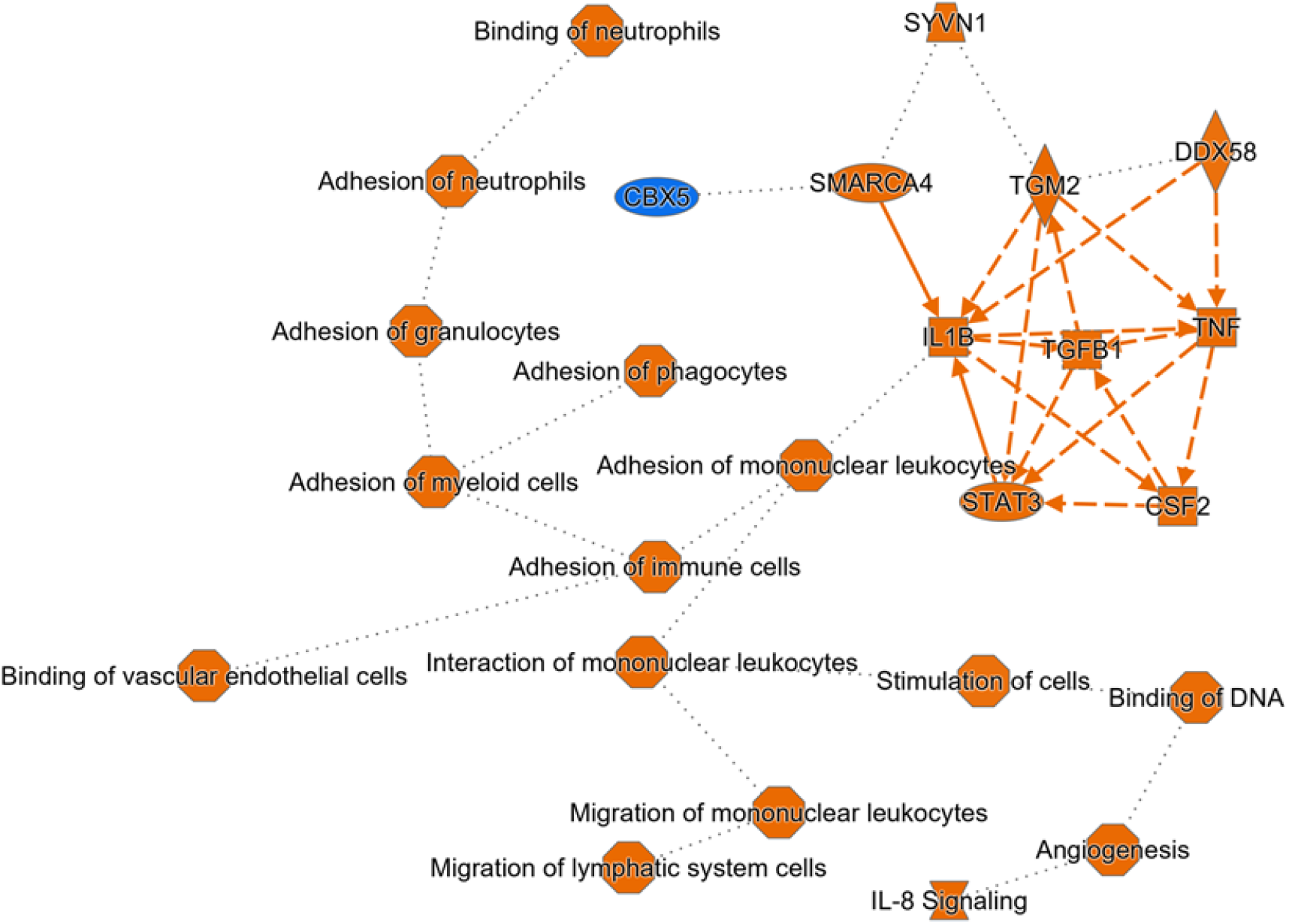
Graphical summary of major biological processes in differentially expressed proteins between controls and NSCLC patients as determined by Ingenuity Pathway Analysis (IPA) software. A solid line indicates a direct interaction, a dash line indicates an indirect interaction, and a dotted line indicates an inferred correlation. Orange symbols indicate activation in NSCLC and blue indicate deactivation.

After applying the filter to the data, a set of 13 DE proteins that were highly predictive of disease status were identified and receiver operating characteristic (ROC) curve analysis revealed a mean area under the ROC curve (AUC) of 0.864 for the set (full list outlined in **Table 3**). All but 3 were elevated in the NSCLC cohort, with an average log fold change of 1.11 ± 0.65 for the increased proteins and - 0.60 ± 0.21 for the decreased proteins.

**Table 3.** Differentially expressed proteins between healthy controls and NSCLC patients after data filtering for a subset of key biomarker candidates.

| Gene | Protein ID | Protein Names | Adjusted p-value | Log fold change | AUC | Sensitivity | Specificity |
| --- | --- | --- | --- | --- | --- | --- | --- |
| SMC2 | O95347 | SMC2 | 0.000131 | -0.85 | 0.972 | 0.94 | 0.94 |
| TP53I11 | O14683 | P5I11 | 0.000217 | 2.30 | 0.931 | 0.88 | 0.78 |
| SIRPA | P78324;<br>Q5TFQ8;<br>O00241 | SHPS1;<br>SIRB1;<br>SIRBL | 0.002688 | 1.30 | 0.885 | 0.75 | 0.83 |
| KRT8 | P05787;<br>P08729;<br>P41219;<br>P17661;<br>P08670 | K2C8;<br>VIME | 0.003865 | 1.92 | 0.885 | 0.81 | 0.78 |
| MACROH2A1 | Q9P0M6;<br>O75367 | H2AW;<br>H2AY | 0.004846 | 1.15 | 0.851 | 0.69 | 0.78 |
| PLP2 | Q04941 | PLP2 | 0.005605 | 0.51 | 0.910 | 0.75 | 0.83 |
| LTF | P02788 | TRFL | 0.006538 | 1.11 | 0.833 | 0.81 | 0.61 |
| UROD | P06132 | DCUP | 0.007004 | -0.54 | 0.861 | 0.81 | 0.78 |
| PLSCR1 | O15162 | PLS1 | 0.008392 | 0.59 | 0.851 | 0.69 | 0.83 |
| CCDC88B | A6NC98 | CC88B | 0.010321 | 1.33 | 0.819 | 0.81 | 0.72 |
| CYB561D2 | O14569 | C56D2 | 0.012271 | 0.61 | 0.823 | 0.75 | 0.78 |
| SEC14L2 | O76054 | S14L2 | 0.048358 | -0.42 | 0.819 | 0.75 | 0.78 |
| SLC27A2 | O14975 | S27A2 | 0.057731 | 0.25 | 0.795 | 0.69 | 0.78 |

Heatmap analysis revealed good separation between NSCLC and healthy controls using this filter (**Figure 6**). A search of Protein Atlas for any known pathology prognostics revealed that 11/13 are associated with various cancers, but none associated with lung cancer. A further manual search of the literature revealed only 2 have any possible known association with lung cancer. These are TP53I11 (tumor protein p53-inducible protein 11) and SLC27A2 (very long-chain acyl-CoA synthetase, ACSVL1).^16,17^

**Figure 6.**
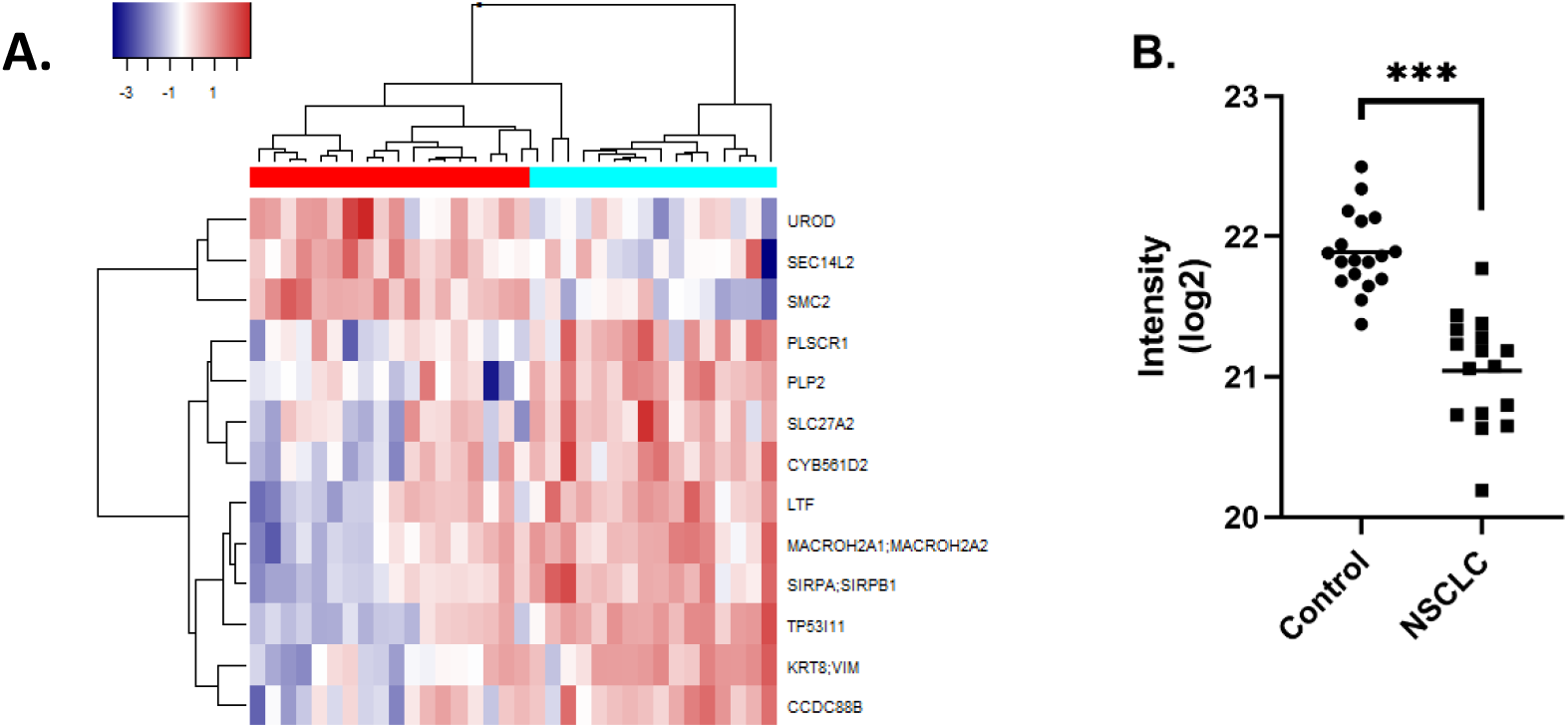
(A) Heatmap of differentially expressed proteins (rows) after parsing though false rate discovery and gradient boosting methods importance filter where samples are listed as columns with controls clustered on the left and NSCLC clustered on the right. (B) Intensity for SMC2 protein in healthy controls and NSCLC patients as measured by mass spectrometry. Data is significant (***) if p < 0.001.

Sensitivity and specificity analysis using logistic regression for each protein revealed one particularly promising protein out of this list. SMC2, a chromosome organisation protein, was found to be significantly lower in the NSCLC group for this cohort (*p* = 0.0001). In classifying NSCLC status, it was found to have a sensitivity of 0.94, a specificity of 0.94 and an AUC of 0.97 (**Figure 6**).

## 5. Discussion

The primary aim of this study was to test a series of novel sample preparation techniques that we have developed for their applicability in identifying disease specific biomarkers. For this purpose, NSCLC was selected as the test cohort and age and sex matched healthy controls were used as the comparator. A collection of NSCLC and healthy control blood samples were purchased from an international biobank and analysed.

Although a huge amount of research has been focussed on identifying NSCLC blood-based biomarkers, none of these markers have yet been brought into diagnostic use.^2^ The vast majority of studies use conventional sample preparation methods, that is, proteomic analysis of serum or plasma. Although analysis of plasma and serum is of course valuable, it only contains a small subset of the potential information available in blood.^9^

For our study, the purchased samples had been processed and stored frozen in the biobank. Only plasma and WB cell pellets were made available to us, thus our fractionation options were limited. Although this was an obstacle, we were able to fractionate those frozen blood samples to produce 4 fractions for analysis: (1) plasma, (2) WB pellet membrane-free fraction, (3) RBC membrane releasate, (4) WB pellet in VAMS.

Using these novel sample preparation methods, 522 DE proteins were identified, with conventional blood analysis (proteomic analysis of plasma) accounting for only 7 of that total. Our methods combined both targeted immunoassays to investigate known low abundance markers as well as discovery mass spectrometry which lacks the sensitivity of immunoassay but is not limited to analysis of specific subsets of known markers. This combined approach in itself is not novel, but the included analysis of RBC fractions and WB cell pellets using these methods are.

The VAMS processing and mass spectrometry analysis methods were based on two earlier publications where they utilised a series of gentle wash steps and in-device protein digestion that effectively removes high abundance proteins thus enabling the detection of a large number of lower abundance proteins from a typically hemoglobin-rich sample.^13,18^ Van Den Broek et al^18^ reported 423 IDs and Molloy et al^13^ reported 1600 protein IDs from their samples. Our improved methods yielded more than both with an average of 3914 protein IDs (**Figure 3**). The potential for biomarker discovery increases with every additional protein ID.

### 5.1. Previously reported biomarkers

To test of the validity of our results, a cross-check of previously identified markers for NSCLC was performed against this dataset. A recent review^19^ summarised a list of the top potential biomarkers for NSCLC which was comprised of 31 proteins. Of those, 12 were detected in our study and 6 of those were found to be DE in the NSCLC cohort, specifically CEACAM, CNDP, VIM, s100A4, COX2, and Her2. This provides confirmation that our results are valid, however, not complete likely due to assay selection and mass spectrometry sensitivity limitations.

As an example, Mazzone et al^4^ report CEA, CYFRA21-1, and HGF as potential biomarker candidates. Of these, we only detected HGF as it was on one of the tested immunoassay panels. The absence of CEA and CYRA21-1 in the mass spectrometry analysis is likely due to the lack of sensitivity, they are typically detected in ng/mL levels.^20^

Despite not detecting CYRRA21-1 and CEA specifically, members of their respective families were detected and found to be differentially expressed. For CYFRA21-1, the related protein cytokeratin-8 (K2C8) was found to be differentially expressed in NSCLC and was one of the key 13 proteins identified after data filtering (**Table 3**). It had an AUC of 0.88 and the log fold-change was 1.88 higher in NSCLC patients. Likewise, several CEA cell adhesion proteins (CEACAM-1, 6, & 8) were all detected and identified as DE proteins. Although not CEA specifically, they are related proteins and have also been implicated in NSCLC pathogenesis and as predictive of treatment outcome.^21–23^ CEACAM-6 in particular has been identified as an important marker of the fatal metastasis in NSCLC, leptomeningeal metastasis.^24,25^

Further, prolactin has been proposed as a biomarker^5^ but was not detected in our dataset. However, Prolactin-Inducible Protein and Prolactin Regulatory Element-Binding Protein were both found in our dataset and determined to be differentially expressed in NSCLC.

Our broad analysis captured both known markers as well as associated proteins of known markers, which supports the validity of our methods and also the validity of the markers themselves. If CEA is known to be involved in NSCLC pathogenesis, it is logical that associated proteins would also be involved and thus also differentially expressed. Here we provide evidence of that in select cases.

### 5.2. Novel biomarkers

In addition to known markers, a number of unknown markers were identified as well. The discovery style proteomics performed in this study has its advantages in that it can identify large number of potentially interesting targets. However, with 522 DE proteins to review, it brings with it a host of challenges. Instead of performing a thorough review of the entire list, the data was filtered to select a top subset of markers (**Table 3**).

All but 2 of these proteins were novel and have never been associated with NSCLC before. One particular protein stands out from the list, SMC2 (also known as hCAP-E). SMC2 had a sensitivity and specificity of 0.94 each, as well as an AUC of 0.97. In context, this result is much higher than other markers have yet been able to achieve. For example, Mazzone et al found the best results by combining 5 markers into a panel and reported a sensitivity of 0.49, a specificity of 0.96, and an AUC 0.76.^4^ It should be noted though, our data is limited by a number of factors. It was a much smaller sample size, the NSCLC participants were a combination of Stage II - Stage IV, and no testing set was assessed. As Mazzone et al demonstrates, efficacy of markers changes with each clinical stage and between training and testing sets.^4^ SMC2 is a subunit of Condensin I and as such is typically intracellularly located. This may explain why it has not been previously identified in plasma/serum analyses. Despite this, a loss of expression of the SMC2 gene has been reported in various gastrointestinal cancers^26^ and is associated with breast cancer risk.^27^ It has even been proposed as a potentially novel cancer therapy target.^28^ Although its association with NSCLC is yet to be validated, it appears to be a very promising target.

Although not a strictly novel biomarker, MIF was also found to be differentially expressed in the NSCLC patients exclusively in the RBC membrane releasate samples (**Figure 1**). Gamez-Pozo et al^29^ has previously identified the prognostic value of MIF in NSCLC, but their analysis required a tissue biopsy which inherently much more invasive than a simple blood draw.

### 5.3. Sex effect

One of the intriguing aspects of the results of this study was the observed sex effect (**Figure 3**-**4**). Females with NSCLC had a biomarker profile that was distinct compared to all of the other groups (female controls, male NSCLC, and male controls). Sex differences have been reported previously in NSCLC pathology and also in response to treatment – females are more likely to have overexpression of KRAS and EGFR in tumours^30^ and immune checkpoint inhibitors are more effective in treating the disease in males.^31^ Notably, tumour mutational burden, a predictor of response to treatment in NSCLC, has been recently found to only be effective for females^32^ and in an analysis of metabolites in dried blood spots, Li et al^33^ identified unique biomarker sets for males and females respectively.

Sirtuins have been implicated in NSCLC^34,35^, however a sex difference in biomarkers such as sirtuins has not yet entered the discussion. Further, C-reactive protein (CRP) was identified as one of the DE proteins for the female cohort exclusively. CRP, a general marker of inflammation, has been recommended as a NSCLC biomarker candidate.^36^ Our dataset, however, suggests that markers such as CRP may only potentially interesting as a biomarker for females with NSCLC, not the entire cohort.

Although this dataset is small, it is indicative that there may be sex biases in proteomic biomarkers of NSCLC as well. A recent publication^6^ hypothesised that multi-protein biomarker panels are more robust than single-protein biomarkers and are more effective at overcoming biological complexities such as sex or lifestyle. Although in this dataset when separated by sex there was a stronger discrimination between controls and cases for the female cohort, the filtered set of 13-biomarkers identified with bootstrapping regression tree analysis was determined to be to highly selective across all participants regardless of sex which supports Baders hypothesis.

## 6. Conclusion

Using a series of novel models, we demonstrate how stored, biobanked samples and be leveraged to produce large reproducible datasets for biomarker discovery. We further provide evidence that there is value to be gained from analysis of additional blood fractions in addition to plasma. Using these methods, a novel 13-protein panel of potential biomarkers of NSCLC was identified that was significant regardless of sex despite a clear sex effect observed in the data.

## 8. Supplementary information

**Table S1.** DIA variable windows used for analysis.

| DIA Window | Centre Mass (m/z) | Start (m/z) | End (m/z) | Width (m/z) |
| --- | --- | --- | --- | --- |
| 1 | 372 | 350 | 394 | 44 |
| 2 | 408.5 | 393 | 424 | 31 |
| 3 | 437.5 | 423 | 452 | 29 |
| 4 | 464.5 | 451 | 478 | 27 |
| 5 | 490.5 | 477 | 504 | 27 |
| 6 | 516 | 503 | 529 | 26 |
| 7 | 541.5 | 528 | 555 | 27 |
| 8 | 567.5 | 554 | 581 | 27 |
| 9 | 594 | 580 | 608 | 28 |
| 10 | 621 | 607 | 635 | 28 |
| 11 | 648.5 | 634 | 663 | 29 |
| 12 | 677.5 | 662 | 693 | 31 |
| 13 | 708.5 | 692 | 725 | 33 |
| 14 | 741.5 | 724 | 759 | 35 |
| 15 | 778 | 758 | 798 | 40 |
| 16 | 819 | 797 | 841 | 44 |
| 17 | 866 | 840 | 892 | 52 |
| 18 | 925 | 891 | 959 | 68 |
| 19 | 1010 | 958 | 1062 | 104 |
| 20 | 1355.5 | 1061 | 1650 | 589 |

**Figure S1.**
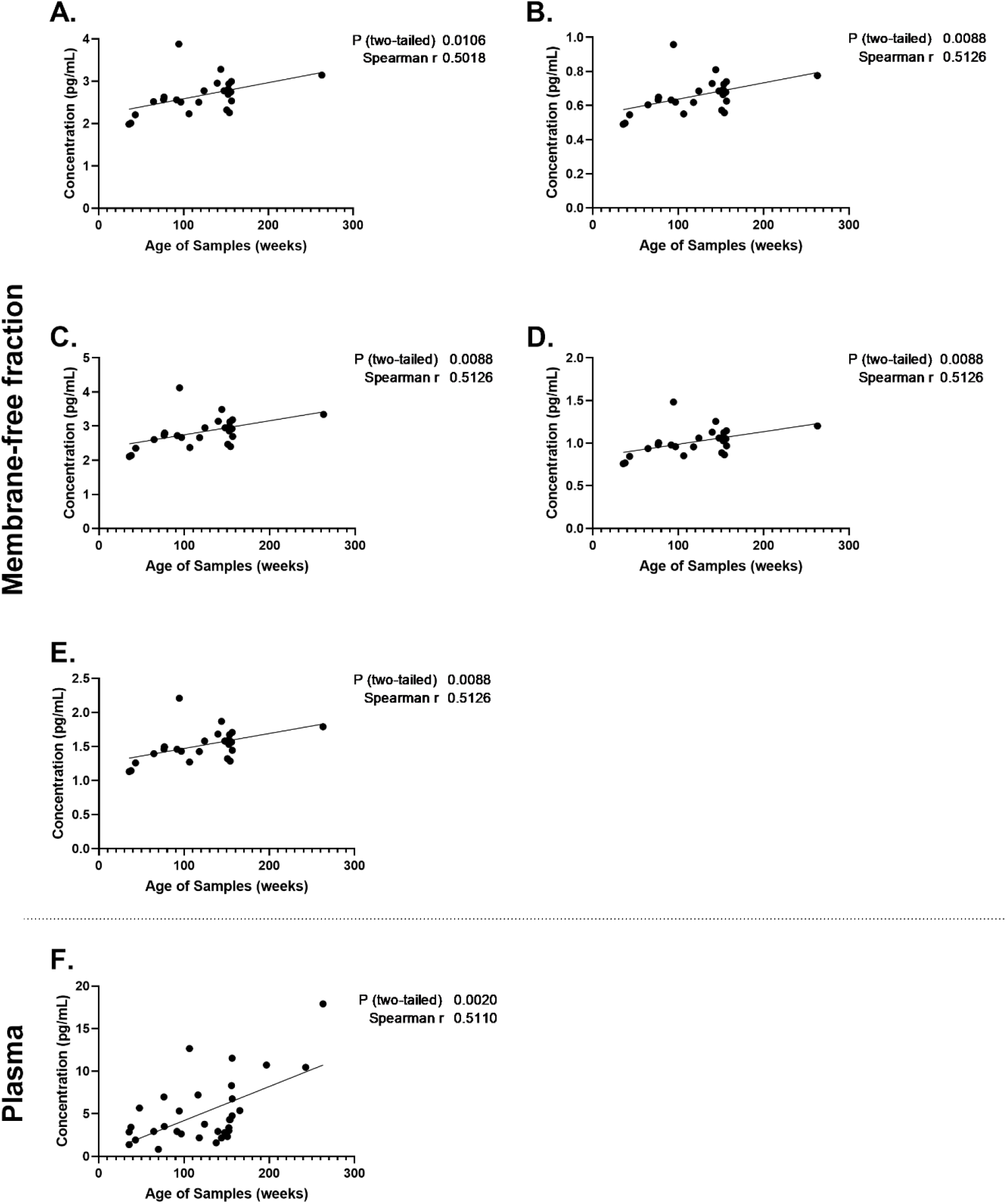
Correlation of (A) Angiopoietin-2, (B) FGF-basic, (C) G-CSF, (D) M-CSF, (E) TGFα from in membrane-free fraction of whole blood cell pellet and (F) Gro-α/KC from plasma with sample age. Correlation determined using Spearman correlation and only analytes with correlation of 0.5 or greater displayed. Data are presented with linear regression line, and p-value and r-value of analysis indicated on each plot. Data deemed significant if p < 0.05.

